# Overcoming Resistance to Anabolic Selective Androgen Receptor Modulator (SARM) Therapy in Experimental Cancer Cachexia with Histone Deacetylase Inhibitor AR-42

**DOI:** 10.1101/214155

**Authors:** Yu-Chou Tseng, Sophia G. Liva, Anees M. Dauki, Michael Sovic, Sally E. Henderson, Yi-Chiu Kuo, Jason A. Benedict, Samuel K. Kulp, Moray Campbell, Tanios Bekaii-Saab, Mitchell A. Phelps, Ching-Shih Chen, Christopher C. Coss

## Abstract

**Purpose:** The common colon-26 mouse (C-26) model of experimental cachexia mimics recent late stage clinical failures of anabolic anti-cachexia therapy, and does not respond to the anabolic selective androgen receptor modulator (SARM) GTx-024. Based on the demonstrated anti-cachectic efficacy of the histone deacetylase inhibitor (HDACi) AR-42 in this model, we hypothesized that combined SARM/AR-42 would provide improved anti-cachectic efficacy.

**Design:** In the C-26 model, we determined a reduced efficacious dose of AR-42 which was combined with anabolic SARM therapy and evaluated for anti-cachectic efficacy. The effects of treatment and tumor burden on anabolic and catabolic signaling occurring in skeletal muscle were characterized using muscle performance parameters and RNA-seq.

**Results:** Anabolic anti-cachexia therapy with diverse androgens had no impact on cachectic outcomes in the C-26 model. A reduced dose of the HDACi AR-42 alone provided limited anti-cachectic benefits, but when combined with the SARM GTx-024, significantly improved bodyweight (p<0.0001), hind limb muscle mass (p<0.05), and voluntary grip strength (p<0.0001) versus tumor-bearing controls. Reduced-dose AR-42 treatment suppressed the IL-6/GP130/STAT3 signaling axis without significantly impacting circulating cytokine levels. GTx-024-mediated β-catenin target gene regulation was apparent in cachectic mice only when combined with AR-42.

**Conclusions:** Cachectic signaling in the C-26 model is comprised of catabolic signaling insensitive to anabolic GTx-024 therapy and a blockade of GTx-024-mediated anabolic signaling. AR-42 treatment mitigates catabolic gene activation and restores anabolic responsiveness to GTx-024. Combining GTx-024, a clinically established anabolic therapy, with a low dose of AR-42, a clinically evaluated HDACi, represents a promising approach to improve anabolic response in cachectic patient populations.

## INTRODUCTION

Cancer cachexia is a multifactorial syndrome characterized by the involuntary loss of muscle mass occurring with or without concurrent losses in adipose tissue. The progressive loss of lean mass associated with cachexia results in decreased quality of life, decreased tolerance of chemotherapy and reduced overall survival [1]. It is estimated that 50-80% of all cancer patients experience cachexia symptoms and up to 20% of all cancer-related deaths are attributable to complications arising from cachexia-mediated functional decline [2]. A multitude of tumor and host factors are recognized as contributors to the multi-organ system dysfunction in cancer cachexia, presenting a considerable therapeutic challenge. Diverse cachexia treatment strategies have been evaluated in patients with few offering effective palliation and none gaining FDA approval for this devastating consequence of advanced malignancy [3]. Among the complex sequelae associated with cachectic progression, compromised muscle function associated with reduced muscle mass is viewed as a primary contributor to patient morbidity and mortality [1]. Recognizing this feature of cancer cachexia, regulatory agencies require the demonstration of meaningful improvements in physical function in addition to improvements in patient body composition for successful registration of novel cachexia therapies [4]. Anabolic androgenic steroids or steroidal androgens are among the most well recognized function-promoting therapies [5] and, as such, have been extensively evaluated in muscle wasting of diverse etiology [6]. Despite meeting FDA approval criteria in other wasting diseases [6], steroidal androgens are yet to demonstrate clinical benefit in cancer cachexia. However, the continued development of novel androgens for the treatment of wasting diseases suggests confidence in this therapeutic strategy remains [3].

In addition to their well characterized anabolic effects on skeletal muscle, steroidal androgens elicit a number of undesirable virilizing side effects and can promote prostatic hypertrophy which limits their widespread clinical use [7]. Recently developed, non-steroidal, selective androgen receptor modulators (SARMs) offer a number of improvements over steroidal androgens including prolonged plasma exposures and oral bioavailability with greatly reduced side effects (virilization, etc.), while maintaining full agonism in anabolic tissues like skeletal muscle [8]. With once daily dosing, the SARM GTx-024 (enobosarm) showed promising gains in fat-free mass in both male and female cancer patients, but ultimately failed to demonstrate a clear functional benefit in pivotal phase III trials in a cachectic non-small cell lung cancer (NSCLC) population [9]. GTx-024 has a strong safety profile and proven effects on skeletal muscle, but is no longer being developed for cancer cachexia.

Hypogonadism is a feature of advanced malignancy and experimental cachexia that worsens multiple cachectic sequelae, including decreased skeletal muscle mass, providing a rationale for therapeutic exogenous androgen administration [10, 11]. Though the relationship between androgen status and body composition is well established, the exact molecular basis by which androgens modulate skeletal muscle mass is not completely characterized but involves the repression of several atrogenes, induction of PI3K/AKT/mTOR signaling and direct stimulation of muscle satellite cells (MUSCs) [12]. Despite demonstrating the clear ability to attenuate orchiectomy- and glucocorticoid-mediated muscle loss [13], in our hands, SARMs displayed essentially no impact on muscle wasting associated with the common colon-26 (C-26) mouse model of experimental cancer cachexia (*unpublished*). In these mice, androgen-mediated gene transcription was severely muted in the presence of a cachectic burden, and fully anabolic doses of SARM were unable to normalize muscle E3-ligase expression to effectively combat the catabolic decline driven by the C-26 tumor. We hypothesized that a better understanding of the failure of SARMs in the C-26 model would offer key insights into the limitations of androgens in cachectic cancer patients. Furthermore, we recently demonstrated the effectiveness of a novel class I/IIB HDAC inhibitor (HDACi, AR-42), currently under clinical evaluation in hematologic malignancies [14] and solid tumors, as anti-cachexia therapy in the C-26 model [15]. AR-42 administration in these mice spared body weight and was associated with improvements, but not complete rescue, of skeletal muscle mass relative to controls. Notably, AR-42 differed from other approved HDACis in its ability to fully suppress tumor-mediated atrogin-1 and MuRF1 induction and prolong survival in the C-26 model. Unlike androgens, AR-42 does not promote skeletal muscle hypertrophy in tumor-free animals, and AR-42’s anti-cachectic efficacy was highly dependent on early initiation of treatment, suggesting AR-42’s dramatic anti-cachectic efficacy in the C-26 model is primarily associated with its anti-catabolic effects [15]. Given SARMs’ established anabolic potential, but clear inability to attenuate tumor-driven wasting in the C-26 model, and AR-42’s effects on tumor-mediated catabolic signaling, but apparent lack of anabolic effects on skeletal muscle, we hypothesized that co-administration of AR-42 with SARMs in the C-26 model would result in improved anabolic response and overall anti-cachectic efficacy. Before evaluating combination therapy, we first explored reduced anti-cachectic doses of AR-42 given the need to minimize side-effects attributable to anti-cachectic therapy.

## MATERIALS AND METHODS

Information on reagents and chemicals, antibodies, cell culture, animals, and methodological details for grip strength measurement, luteinizing hormone analysis, pharmacokinetics, western blot analysis and gene expression analyses are included in the *Supplementary Materials and Methods*. All animal studies were conducted according to protocols approved by The Ohio State University Institutional Animal Care and Use Committee.

### Animal studies using the C-26 colon adenocarcinoma cachexia model

These studies were performed as previously described [15] with modifications. Tumors were established in the right flank by subcutaneous injection of C-26 cells (0.5 × 10^6^ cells in 0.1 mL). AR-42, GTx-024 and their vehicles were administered orally by gavage. TFM-4AS-1 (a potent experimental SARM), dihydrotestosterone (DHT) and vehicles were administered by subcutaneous injection.

#### AR-42 dose-response study

Male CD2F1 mice were stratified by body weight and then randomly assigned into 5 groups of 6 animals each. C-26 tumors were established in four of the groups, while those in the fifth group, serving as tumor-free controls, were injected with sterile saline. Six days later, animals with palpable tumors were treated with AR-42 once daily at 10 (n=5) and 20 mg/kg (n=6), and every other day at 50 mg/kg (n=5), or vehicle control (n=4) for 13 days. Upon sacrifice on study day 18, when the majority of tumor-bearing control mice met euthanasia criteria, the left gastrocnemius muscle was excised, flash frozen in liquid nitrogen and stored at −80°C for subsequent analyses. Carcass weights were corrected for tumor weight by assuming a tumor density equivalent to water (1 g/cm^3^).

#### Initial AR-42/GTx-024 Combination Study (Study 1)

Male CD2F1 mice were stratified by body weight and then randomly assigned into 6 groups of 6 animals each. Historically, 6 animals per group provided sufficient power to detect treatment-mediated differences in tumor-bearing treated animals compared to controls. Tumors were established in four of the groups, while the fifth and sixth groups served as tumor-free controls. Six days later, animals with palpable tumors were treated twice daily for 13 days. AR-42 and its vehicle were administered in the mornings, and GTx-024 and its vehicle in the afternoons. Treatments included vehicles for AR-42 and GTx-024 (n=5), GTx-024 (15 mg/kg; AR-42 vehicle; n=5), AR-42 (10 mg/kg; GTx-024 vehicle; n=5), or the AR-42+GTx-024 combination (10 and 15 mg/kg, respectively; n=5). The remaining tumor-free groups received either vehicles (n=6) or GTx-024 (15 mg/kg; AR-42 vehicle; n=6). Body weight, tumor volume, and feed consumption were monitored every other day. Upon sacrifice on day 18, sera were collected and hind limb skeletal muscles, heart, spleen and epididymal adipose tissues were harvested, weighed, flash frozen, and stored for subsequent analyses.

#### Confirmatory AR-42/GTx-024 Combination Study (Study 2)

This confirmatory study was performed exactly as Study 1 with expanded animal numbers. Tumor-free control groups were maintained at 6 animals each, whereas 10 animals were included in each of the tumor-bearing groups. Six days after cell injection, animals with palpable tumors were treated as in Study 1 with vehicles (n=7), GTx-024 (n=10), AR-42 (n=9), or the combination (n=9). Grip strength was measured on study days 0 (baseline) and 16. Due to rapid model progression, this study was terminated after only 12 days of treatment.

#### Combined Androgen and AR-42 Study (Study 3)

Similar to Study 2, the tumor-free control group was maintained at 6 animals, whereas 10 animals were included in each of the 6 tumor-bearing groups. Six days after cell injection, animals with palpable tumors were treated once daily for 13 days with vehicles for AR-42 and TFM-4AS-1/DHT (n=9), AR-42 (10 mg/kg; TFM-4AS-1/DHT vehicle; n=10), TFM-4AS-1 (10 mg/kg; AR-42 vehicle; n=9), DHT (3 mg/kg; AR-42 vehicle; n=10), the combination of AR-42 and TFM-4AS-1 (10 mg/kg each; n=9), or the combination of AR-42 (10 mg/kg) and DHT (3 mg/kg; n=10). Grip strength was measured and tissues collected as in the previous studies.

### AR-42 Plasma and Tissue Pharmacokinetics

Pharmacokinetic studies were performed as previously described [16] with modifications described in *Supplementary Materials and Methods*.

### Western Blot Analyses

All Western blots were performed on gastrocnemius muscle from representative animals in each study using standard methods as described in *Supplementary Materials and Methods*.

### Gene Expression Analyses

All gene expression experiments were performed on gastrocnemius muscles from representative animals. Expression levels were estimated by both traditional q-RT-PCR and sequencing of polyA-selected RNAseq libraries. Details of the methods for data generation and associated analyses are described in detail in *Supplementary Materials and Methods*.

### Statistical Methodology

Plotting and statistical analyses were performed using GraphPad Prism Version 7 (GraphPad Software, La Jolla, CA). The specific statistical tests employed are outlined in detail within the figure legends.

## RESULTS

### AR-42 administration demonstrates anti-cachectic effects at a reduced 10 mg/kg dose level

We recently characterized the anti-cachectic effects of AR-42 (50 mg/kg via oral gavage every other day) in C-26 tumor-bearing mice [15]. This dose represented the maximally tolerated dose in mice, which was used to observe its anti-tumor effects in different xenograft tumor models. To better understand the disposition of AR-42 following oral administration in mice, we performed a limited pharmacokinetic study of single oral doses of 50, 20 and 10 mg/kg of AR-42 (Figure 1A). Plasma exposure following oral administration of 50 mg/kg was 74.3 µM*h (Supplementary Figure S1), which exceeded the well tolerated plasma exposure in humans of 8.5 µM*h by 8.7-fold [14]. Consequently, we evaluated the anti-cachectic effects of lower doses of AR-42 in a dose-response study in the C-26 model. Similar to six total 50 mg/kg doses (administered q2d), thirteen daily oral doses of 20 or 10 mg/kg AR-42 reversed C-26 tumor-mediated reductions in tumor-corrected body weight (Figure 1B). AR-42 readily distributed into gastrocnemius muscle tissue (Figure 1A) and, at the lowest efficacious dose of 10 mg/kg, muscle concentrations remained above 700 nM for 4 hours consistent with the ability of AR-42 at this dose to inhibit Class I and IIb HDACs for a portion of the dosing interval in muscle tissue based on its *in vitro* HDAC inhibition profile (Figure 1C). The plasma exposure resulting from the 10 mg/kg dose (10.9 µM*h, Supplementary Figure S1) compares more favorably to well-tolerated exposures in patients while providing anti-cachectic efficacy and was therefore utilized in subsequent combination studies.

**Figure 1.**
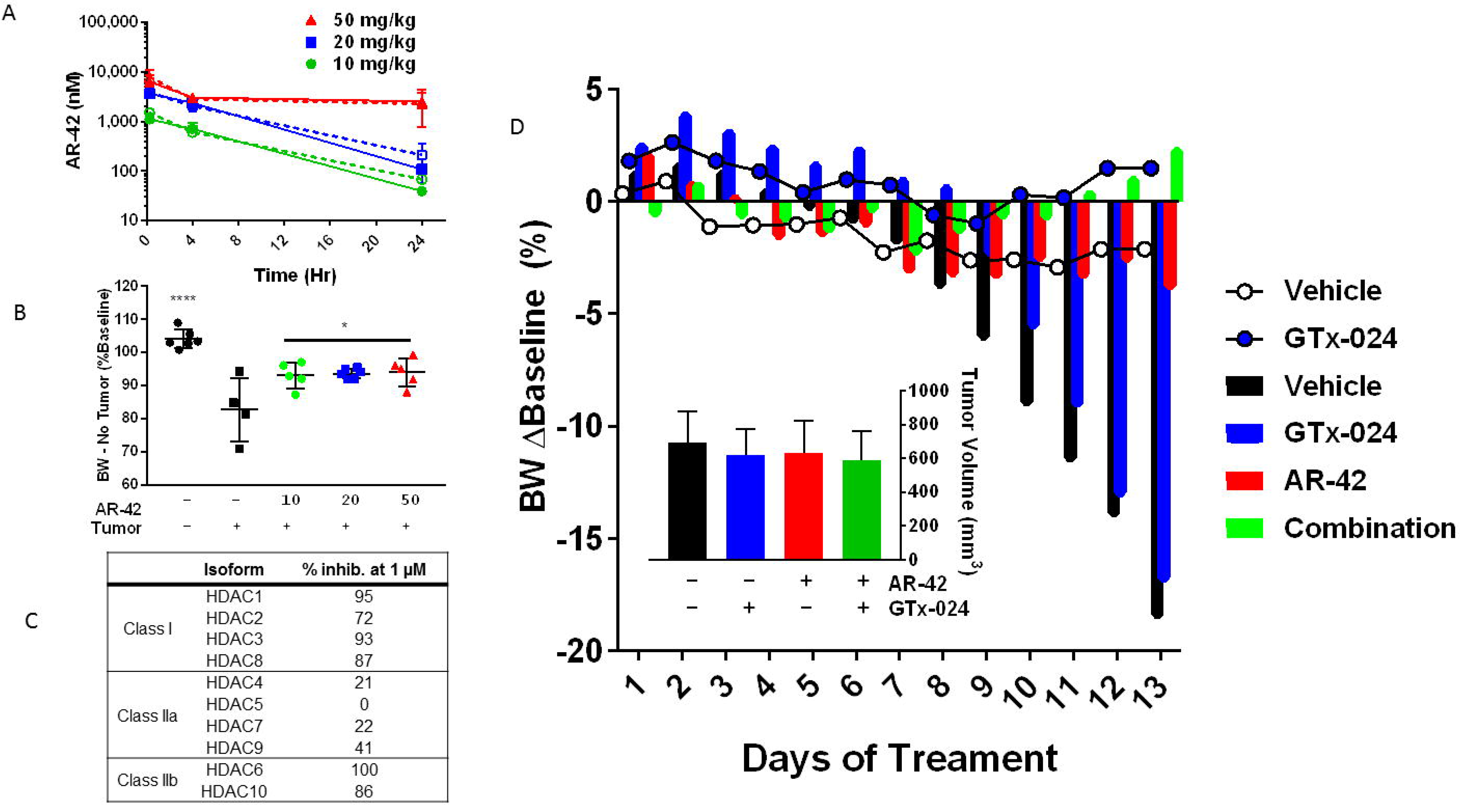
**A)** Single dose AR-42 plasma and tissue pharmacokinetic study. Tumor-free C57BL/6 mice were administered a single dose of 10, 20 or 50 mg/kg AR-42 (n=3) and plasma (dashed) and gastrocnemius (solid) tissue were analyzed for AR-42 content at different times using LC-MS/MS analyses according to *Materials and Methods* (Mean±SD). **B)** AR-42 dose-response. Starting six days after C-26 cell injection, animals received vehicle or AR-42 orally at 10 or 20 mg/kg daily or 50 mg/kg every other day for 13 days (n=4-6). Terminal (Day 18 post-injection) body weights compared to baseline (Day 0), corrected for tumor mass according to the *Materials and Methods*. Bars represent mean±SD. *p<0.05, ****p<0.0001 versus tumor-bearing vehicle-treated controls, Tukey’s multiple comparison test. **C)** AR-42’s *in vitro* human HDAC inhibition profile was determined using recombinant enzymatic assays according to *Materials and Methods*. **D)** Study 1, Animals receiving GTx-024 (15 mg/kg), AR-42 (10 mg/kg), Combination (15 mg/kg GTx-024 and 10 mg/kg AR-42) or Vehicle were treated daily by oral gavage for 13 days starting 6 days post-injection of C-26 cells. Longitudinal mean body weights per treatment group are presented as a percent change from pre-cell injection body weights. Tumor-free animals (circles), tumor-bearing animals (bars). *Inset:* terminal tumor volumes (mean±SD, n=5-6 per group).

### Combination GTx-024 and AR-42 administration results in improved anti-cachectic efficacy

To evaluate our hypothesis that combining HDAC inhibition with SARM administration would improve anti-cachectic activity, we designed a series of three studies combining AR-42 with androgen/SARM in the C-26 model. In Study 1, similar to results from this and other laboratories [15, 17], vehicle-treated tumor-bearing animals lost approximately 20% of their body weight prior to meeting euthanasia criteria (Figure 1D). This severe tumor-induced weight loss (Figure 2A, 80.4±9.1% of baseline) was accompanied by parallel reductions in gastrocnemius and quadriceps masses (Figure 2B, 86±12.4 and 88±12.0%, relative to tumor-free controls, respectively). Consistent with previous findings in this model (*unpublished*), SARM monotherapy had no apparent anti-cachectic efficacy in C-26 tumor-bearing mice. GTx-024 at 15 mg/kg did not spare body weight (Figure 1D, 2A) or the mass of gastrocnemius and quadriceps muscles (Figure 2B). At this dose, GTx-024 was well tolerated in xenografted mice [18] and, in this study, did not cause body weight loss in tumor-free controls (Figure 1D, 2A).

**Figure 2.**
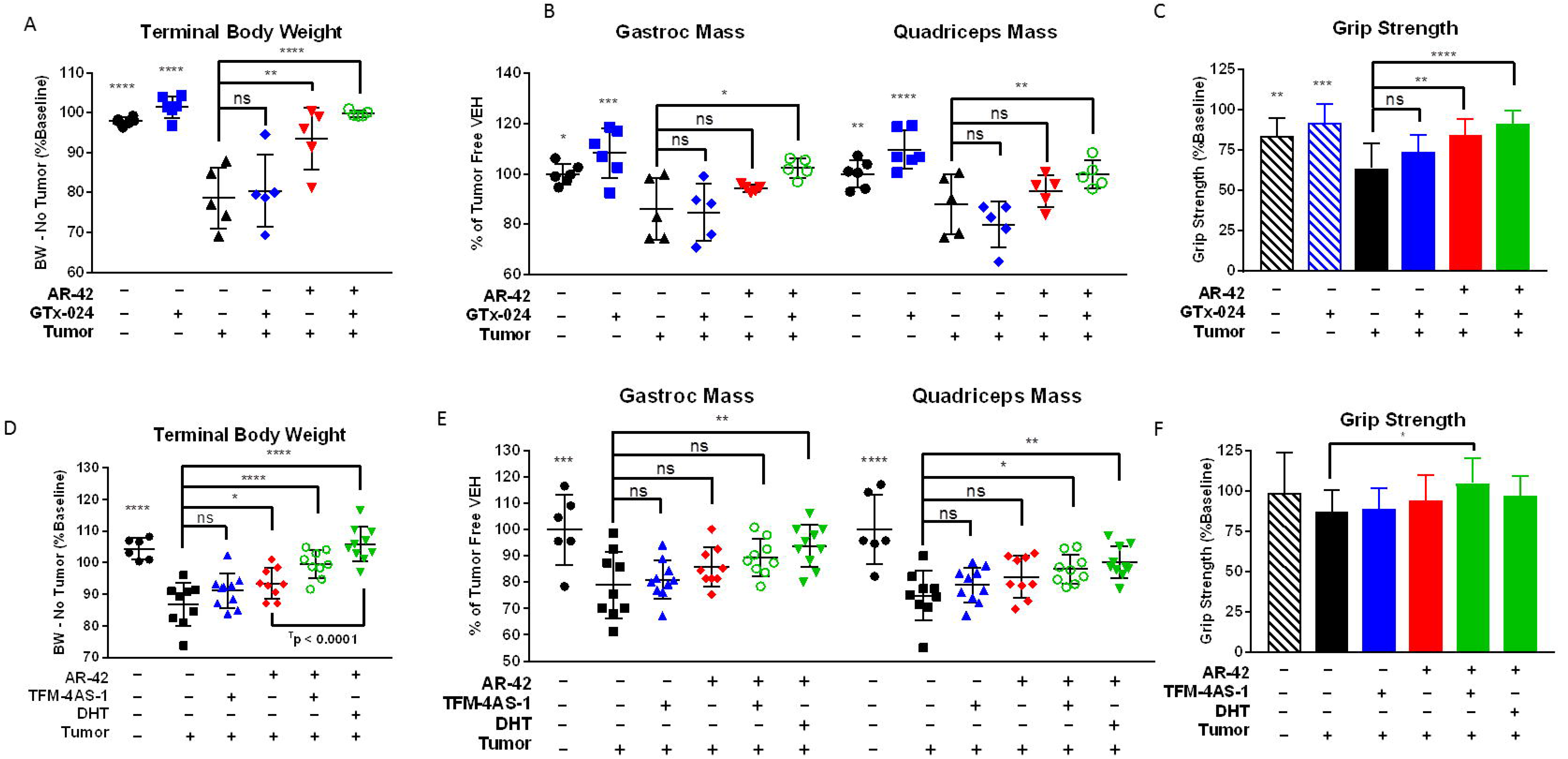
**A-B)** Study 1, Animals receiving GTx-024 (15 mg/kg), AR-42 (10 mg/kg), Combination (15 mg/kg GTx-024 and 10 mg/kg AR-42) or Vehicle were treated daily by oral gavage for 13 days starting 6 days post-injection of C-26 cells (n=5-6 per group). **A)** Terminal (Day 18 post-injection) body weights compared to baseline (Day 0), corrected for tumor mass according to the *Materials and Methods*. **B)** Terminal hindlimb skeletal muscle masses. **C)** Day 16 grip strength measurements are presented from Study 2 (n=6-10 per group) compared to pre-treatment baseline. **D-F)** Study 3, Animals receiving AR-42 (10 mg/kg, oral gavage), TFM-4AS-1 (10 mg/kg, subcutaneous), Combination AR-42 and DHT (10 mg/kg oral gavage and 3 mg/kg subcutaneous, respectively), Combination AR-42 and TFM-4AS-1 (10 mg/kg, both) or Vehicle were treated daily for 12 days starting 6 days after cell injection (6-10 per group). **D)** Terminal (Day 17) body weights compared to baseline (Day 0), corrected for tumor mass according to the *Materials and Methods*. **E)** Terminal hindlimb skeletal muscle masses. **F)** Grip strength measurements performed on the final day of treatment compared to pre-treatment baseline. *Statistics for all panels*: Mean±SD. *p<0.05, **p<0.01, ***p<0.001, ****p<0.0001 versus tumor-bearing vehicle-treated controls, Dunnett’s multiple comparison test; ns, no significant difference. An additional Tukey’s multiple comparison test was used in panel **2D** to demonstrate AR-42 combined with DHT resulted in significant improvement (p<0.0001) in terminal body weight compared to AR-42 treatment alone.

Furthermore, GTx-024 was reported to be fully anabolic at doses as low as 0.5 mg/kg/day in rodents (reported as S-22 in Kim *et al.* [19]) and compared favorably to the less potent structural analog S-23 [20], which reversed orchiectomy- and glucocorticoid-mediated wasting [13]. A separate control study in tumor-free CD2F1 mice confirmed that, in our hands, 15 mg/kg GTx-024 was capable of increasing body weight, gastrocnemius and quadriceps mass, and grip strength in orchiectomized (ORX) mice relative to vehicle-treated ORX controls (Supplementary Figure S2). Importantly, GTx-024 suppressed serum luteinizing hormone, a very well characterized pharmacological effect of potent androgen administration [21], demonstrating that GTx-024 administered to C-26 tumor-bearing mice was active (Supplementary Figure S3A).

Consistent with the preliminary dose-response study, 10 mg/kg AR-42 alone significantly spared body weight (Figure 1D and 2A, 93.6±7.7% of baseline) relative to tumor-bearing vehicle-treated controls. However, these changes were not translated into significant improvements in gastrocnemius and quadriceps mass (Figure 2B). In contrast to monotherapy, C-26 tumor-bearing mice receiving both GTx-024 and AR-42 started to gain body weight relative to baseline after nearly two weeks of treatment, whereas all other treated tumor-bearing groups lost body weight (Figure 1D). This combination exhibited a striking ability to consistently protect body weight (99.9±0.8% of baseline, corrected for tumor weight) relative to either agent alone (Figure 2A). Furthermore, the effects of combined therapy completely spared gastrocnemius (102.4±3.8%) and quadriceps (99.9±5.5%) mass relative to tumor-free controls (Figure 2B).

The effects of combined therapy on total body weight or amelioration of cachectic symptoms were not due to any overt impact on tumor burden as no significant differences in tumor volumes were apparent at the end of the study (Figure 1D, inset). Food consumption was monitored to account for potential anti-anorexic effects of treatment on the cachectic sequela following C-26 cell inoculation. GTx-024-treated tumor-free control animals, as well as the combination-treated group, demonstrated small increases in per animal food consumption relative to other groups between day 14 and 16 (Supplementary Figure S3B), which are unlikely to account for differences in body weight apparent by study day 14 (treatment day 9), as well as end of study differences in skeletal muscle masses (Figure 2B).

These promising results prompted us to repeat the experiment with expanded animal numbers and the use of forelimb grip dynamometry as a measure of muscle function. In this confirmatory study (Study 2), the model was more aggressive resulting from significantly larger tumors relative to Study 1, though no differences within treatment groups were apparent (Supplementary Figure S4A). As a result, the study was terminated early, after only 12 days of treatment. In accordance with this increased tumor burden, tumor-corrected body weights were more consistently reduced and to a larger degree in tumor-bearing controls (77.0±5.7% and 80.4± 9.1% of baseline in the second and first studies, respectively; Supplementary Figure S4B), and larger losses in gastrocnemius (76.3±8.1%) and quadriceps (69.1±7.6%) mass relative to tumor-free controls were noted (Supplementary Figure S4C). In the face of this more severe cachexia, only combined AR-42 and GTx-024 administration significantly spared body weight (90.3±4.3 of baseline), though not to the degree realized in the first study (Supplementary Figure S4B vs Figure 2A), while both AR-42 alone and the combination significantly spared gastrocnemius and quadriceps mass (Supplementary Figure S4C). C-26 tumors were accompanied by large reductions in forelimb grip strength (Figure 2C, 63.8±15.3% versus 83.8±10.7% of baseline in tumor-bearing and tumor-free controls, respectively), but, consistent with the improvements in hind limb skeletal muscle mass, AR-42 alone and in combination with GTx-024 improved grip strength over vehicle-treated tumor-bearing controls. Unlike the adipose-sparing effect of the higher 50 mg/kg dose of AR-42 [15], the lower dose of 10 mg/kg had no impact on adipose or heart mass (Supplementary Figure S4D). As androgens are thought to actively prevent adipogenesis [22], SARM administration was not expected to protect against C-26 tumor-mediated fat losses. Indeed, no treatment mediated effects on abdominal adipose were apparent (Supplementary Figure S4D). The data show heart mass was significantly improved by combination therapy, but this result is likely due to the effects of a single outlier animal.

### Multiple androgens demonstrate improved anti-cachectic efficacy when combined with AR-42

To confirm that the improvement of GTx-024’s anti-cachectic efficacy in the C-26 model by co-administration with AR-42 was not a drug-specific phenomenon, tumor-bearing animals were treated with the SARM TFM-4AS-1 [23] and the potent endogenous androgen DHT alone and in combination with AR-42 (Study 3). Similar to the 15 mg/kg dose of GTx-024, TFM-4AS-1 was administered at a previously characterized fully anabolic dose (10 mg/kg), but, as a monotherapy, did not spare body weight (Figure 2D) or mass of gastrocnemius or quadriceps (Figure 2E).

AR-42 alone resulted in significant attenuation of body weight loss (93.5±4.8 of baseline), but was less effective than in combination with TFM-4AS-1 (99.5±4.4 of baseline) or DHT (106.0±5.4 of baseline). The DHT/AR-42 combination significantly improved bodyweights (p<0.0001) compared to AR-42 treatment alone (Figure 2D). Of note, tumor-bearing animals treated with DHT alone did not differ in initial tumor volumes (Day 8), but after 8 days of DHT administration, tumor growth was significantly suppressed resulting in the exclusion of DHT alone treated animals from further analyses (Supplementary Figure S5A). Consistent with both Studies 1 and 2, improvements in body weight were not due to sparing adipose tissue as no treatment-mediated effects on adipose were apparent (Supplementary Figure S5B).

Similar to the first study, AR-42 monotherapy did not significantly impact skeletal muscle masses despite positive effects on body weight. However, combination treatment-mediated improvements in body weight were again translated to increased skeletal muscle masses where DHT/AR-42 combination significantly spared both gastrocnemius and quadriceps mass (93.7±8.0 and 87.5±6.1% versus tumor-free controls, respectively), while the TFM-4AS-1/AR-42 combination attenuated atrophy of the quadriceps only (85 ±5.5% of tumor-free controls, Figure 2E). Congruent with the lesser impact of the C-26 tumors on lower limb skeletal muscle mass in Study 3, smaller deficits in grip strength were apparent in tumor-bearing controls relative to Study 2 (86.5% and 63.8% of baseline, respectively; Figure 2F vs 2C). The only treatment resulting in significantly improved grip strength was the combination of TFM-4AS-1 and AR-42, which increased muscle function over baseline (104.2%) despite the presence of C-26 tumors.

### Effects of tumor burden and GTx-024/AR-42 treatment on the expression of AR and atrophy-related genes in skeletal muscle

Candidate gene expression analyses were performed on gastrocnemius tissue from Study 1 to characterize the effects of C-26 tumors and treatment with GTx-024, AR-42 or both agents on genes whose function has been previously associated with C-26 tumor-mediated wasting (Figure 3A). As expected for this model, the muscle-specific E3 ligases atrogin-1 (FBXO32) and MuRF-1(TRIM63) were induced in skeletal muscles of tumor-bearing animals [15, 17] as was the STAT3 target gene and regulator of atrogin-1 and MuRF-1, CEBPδ(CEBPD) [24]. Consistent with the absence of any anti-cachectic effects of GTx-024 monotherapy, this treatment had no significant impact on atrogin-1, MuRF-1, or CEBPδ expression. Ten mg/kg AR-42 alone and in combination with GTx-024 significantly reduced the expression of each atrogene relative to tumor-bearing controls, returning them to near baseline levels. AR-42’s effects on E3 ligase expression were consistent with results from animals receiving the higher dose of 50 mg/kg [15] further supporting the importance of AR-42’s ability to reverse induction of these key enzymes to its overall anti-cachectic efficacy.

**Figure 3.**
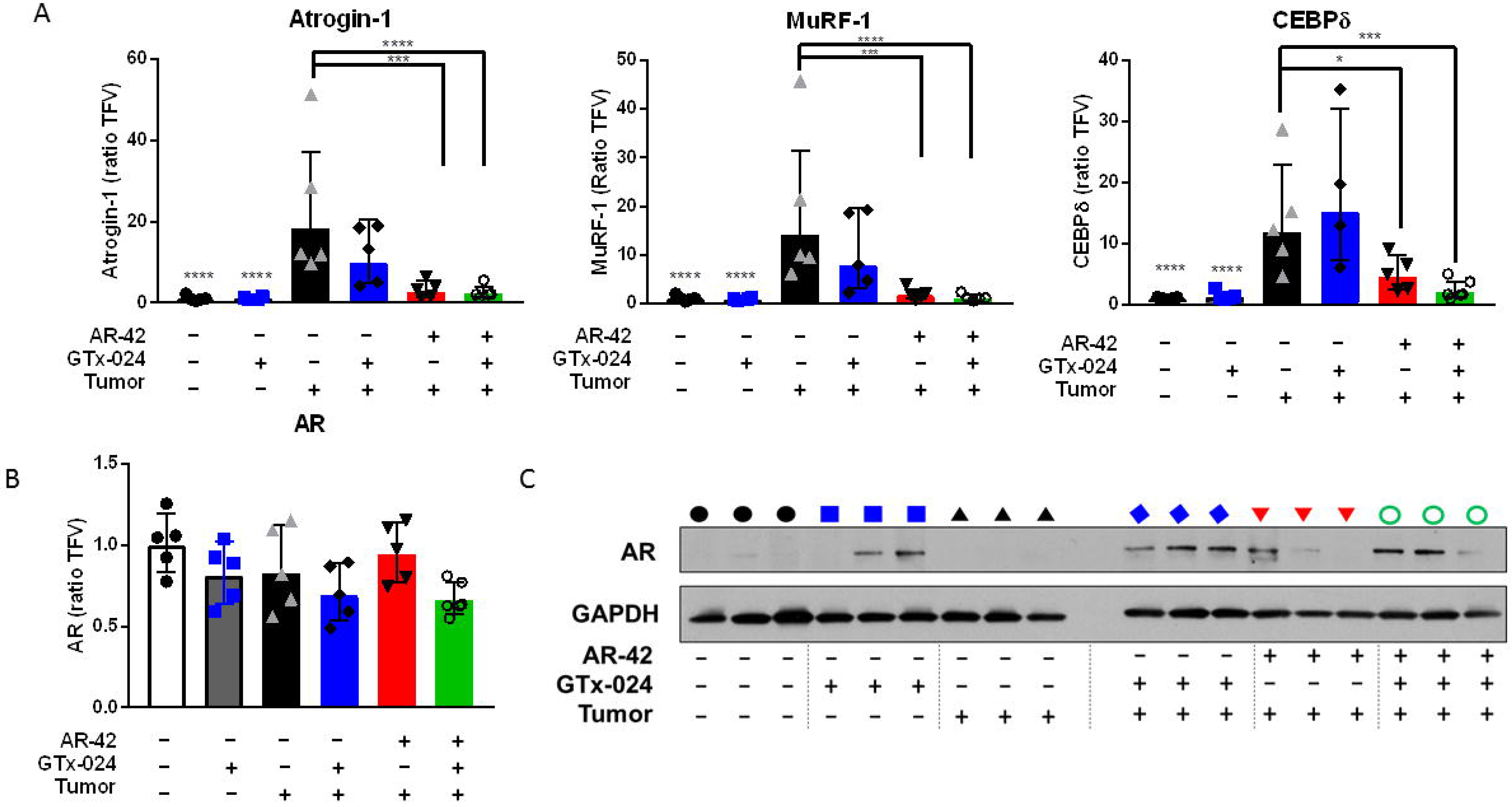
**A-B)** Gene expression of multiple cachexia-associated markers in gastrocnemius muscles of individual animals from Study 1 (n=5-6 per group). Expression was determined by qRT-PCR and presented as described in the *Materials and Methods* (Geometric Mean ± Geometric STD). **A)** Genes associated with muscle atrophy. **B)** Androgen receptor (AR) mRNA expression. **C)** Western blot analysis of AR in gastrocnemius muscles from representative mice in Study 1. *Statistics all panels*: *p<0.05, ***p<0.001, ****p<0.0001 versus tumor-bearing vehicle-treated controls, Dunnett’s multiple comparison test. CEBPδ, n=4, insufficient sample to analyze all tumor-bearing GTx-024-treated animals.

To determine the effect of tumor burden and treatment on androgen receptor (AR) levels in skeletal muscle, which could influence response to androgen therapy, gastrocnemius AR levels were characterized. Neither tumor nor treatment had a significant impact on AR mRNA (Figure 3B). AR protein expression in gastrocnemius was low in tumor-free controls and increased in response to GTx-024 administration irrespective of tumor burden (Figure 3C) consistent with androgen agonist binding and stabilization of the AR [25]. In contrast, AR-42 treatment did not have a marked impact on AR expression.

### Anti-cachectic efficacy of AR-42 is associated with STAT3 inhibition but not general immune suppression

Previously reported ingenuity pathway analyses of AR-42-regulated genes in gastrocnemius muscle revealed that 66 genes associated with muscle disease or function were significantly regulated by AR-42 relative to C-26 tumor-bearing vehicle-treated controls [15]. In an effort to enrich previously reported differentially regulated genes (n=548) for transcripts critical to the anti-cachectic efficacy of AR-42, these data were intersected with previously published differentially regulated genes from the quadriceps of moderate and severely wasted C-26 tumor-bearing mice [17] (n=700, Supplementary Figure S6A). Using this approach, the likely biological relevance of the 147 overlapping genes is increased when it is considered that these transcripts represent genes regulated by AR-42 that are associated with C-26-induced wasting from two different muscles (gastrocnemius and quadriceps), detected by two different technologies (RNA-seq and microarray) and reported by two different research laboratories. Pathway analyses performed on this pool of 147 genes revealed IL-6 signaling and immune system pathways, along with other gene sets regulated subsequent to cytokine stimulation, implicating AR-42’s effects on cytokine and immune signaling in its anti-cachectic efficacy (Supplementary Figure S6B).

In agreement with the present pathway analyses, we previously reported that the higher 50 mg/kg dose of AR-42 reduced serum IL-6 levels, as well as gastrocnemius IL-6 receptor mRNA abundance in tumor-bearing mice suggesting AR-42’s efficacy may be related to its suppression of systemic IL-6 activation which is thought to drive muscle wasting in the C-26 model [15]. In this study, the impact of C-26 tumor burden and treatment with AR-42, GTx-024 or combination therapy on a panel of circulating cytokines, including IL-6, was assessed (Figure 4A, Supplementary Table S1). Consistent with our previous report, multiple pro-cachectic factors, including G-CSF, IL-6, and LIF, were significantly elevated by the presence of C-26 tumors [15]. Unlike the 50 mg/kg dose, 10 mg/kg AR-42 did not significantly impact IL-6 family cytokine levels (i.e. IL-6 or LIF) alone or in combination with GTx-024. Furthermore, 10 mg/kg AR-42 monotherapy did not significantly reduce circulating levels of any evaluated cytokine, despite demonstrating clear anti-cachectic effects across the multiple studies presented here (Figure 1D, Figure 2, Supplementary Figure S4). An ELISA analysis confirmed our findings that AR-42 treatment did not affect circulating IL-6 levels (Figure 4B), and demonstrated serum IL-6 levels were not associated with body weight in treated, C-26 tumor-bearing mice at sacrifice (Figure 4C).

**Figure 4.**
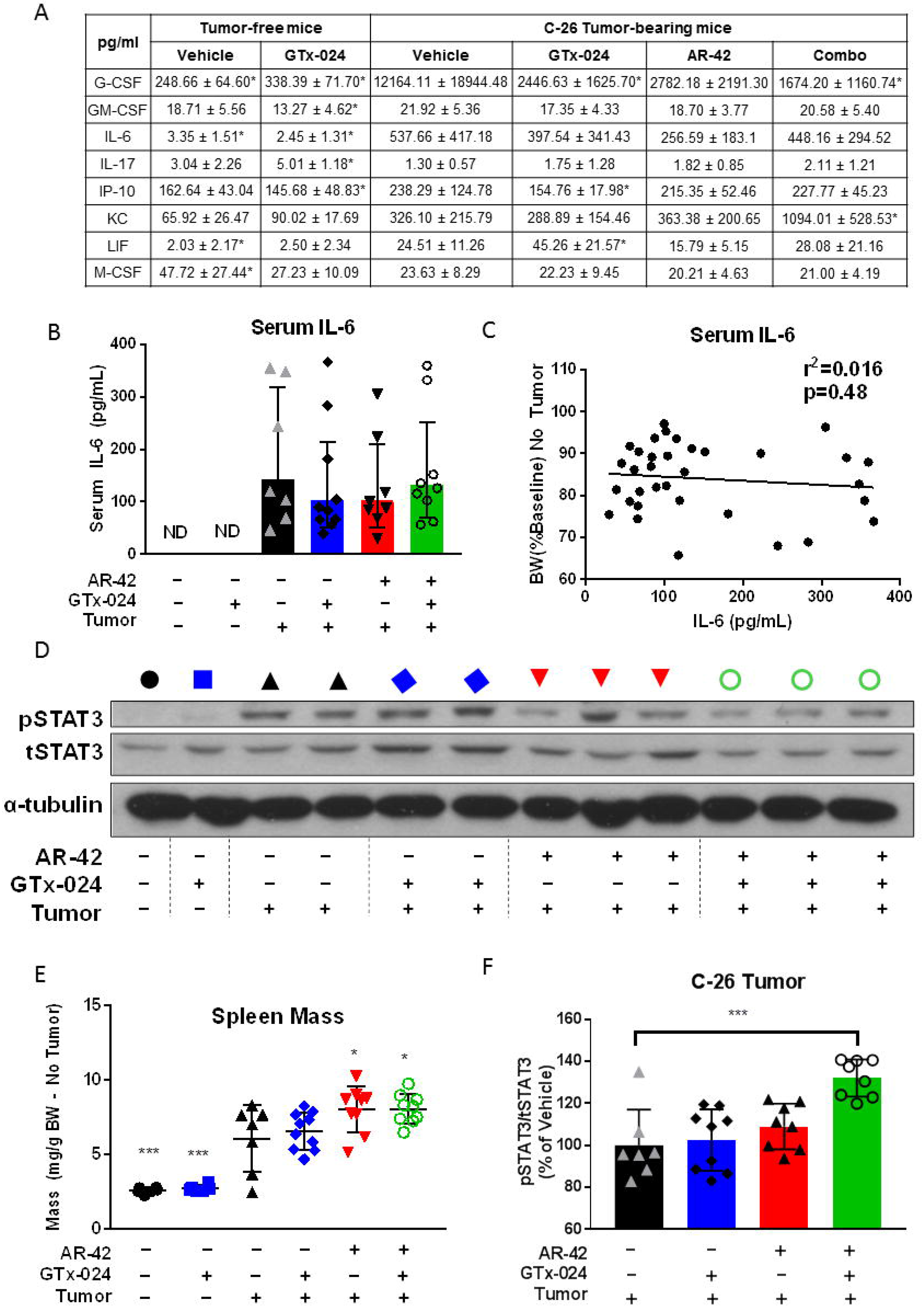
**A)** Multiplex analysis of diverse serum cytokines in terminal samples from Study 2. Presented cytokines are limited to those showing significant differences from tumor-bearing vehicle-treated controls. G-CSF: granulocyte colony-stimulating factor, GM-CSF: granulocyte macrophage colony-stimulating factor, IL-6: interleukin-6, IL-17: interleukin-17, IP-10: interferon gamma-induced protein 10, KC: chemokine (C-X-C motif) ligand 1, LIF: leukemia inhibitory factor, M-CSF: macrophage colony-stimulating factor. Complete cytokine data are presented in Supplementary Table S1. **B)** ELISA analysis of serum IL-6 levels in terminal samples from Study 2. ND, not detected. **C)** Individual animal serum IL-6 values as determined in **A** plotted against tumor-corrected terminal body weights from Study 2 (Pearson’s Correlation). **D)** Phospho(p)STAT3/STAT3 western blot analysis of gastrocnemius tissues from representative animals treated in Study 1. **E)** Spleen weights normalized to tumor-corrected terminal body weights of mice from Study 2. **F)** ELISA analysis of STAT3 within C-26 tumors from Study 2. *Statistics* for *all panels*: Mean±SD, *p<0.05, **p<0.01, ***p<0.001, ****p<0.0001 versus tumor-bearing vehicle-treated controls, Dunnett’s multiple comparison test. Panel **F**, one sample from each of the treatment groups was not available for analyses.

When significant effects on circulating cytokines were not apparent, we hypothesized AR-42 might be acting downstream of the IL-6 receptor on critical mediators of cytokine signaling. One well characterized effector of cytokine-induced signaling shown to be central to tumor-induced wasting in a number of models is signal transducer and activator of transcription (STAT)3 [11, 17]. Notably, STAT3 activation is associated with the severity of wasting in both the C-26 and *Apc*^*min*/+^ models of cancer cachexia, and AR-42 was previously shown to suppress the IL-6/GP130/STAT3 signaling axis in multiple myeloma cells [26]. Thus, we evaluated AR-42’s effects on phospho-STAT3 (pSTAT3) in gastrocnemius muscle from C-26 tumor-bearing animals (Figure 4D, Supplementary Figure S7). As expected, the presence of the C-26 tumor resulted in increased pSTAT3 abundance. GTx-024 treatment had no apparent effect on pSTAT3, consistent with its inability to spare body weight or lower limb skeletal muscle mass as a monotherapy. AR-42 monotherapy reduced pSTAT3 but not equally in all animals, whereas the combination treatment exhibited the most consistent suppression, concordant with its marked anti-cachectic efficacy. Furthermore, treatment-mediated effects on the well characterized STAT3 target gene CEBPδ [24] closely paralleled those on STAT3 activation (Figure 3A).

In addition to skeletal muscle STAT3 activation, C-26 tumor-bearing mice exhibit splenomegaly as a result of increased systemic inflammation [27]. Consistent with increased circulating cytokine levels, C-26 tumor-bearing animals in both Study 1 and 2 demonstrated large increases in spleen mass across all treatment groups relative to tumor-free controls (Supplementary Figure S3C and Figure 4E, respectively). Similar to findings with 50 mg/kg AR-42 [15], spleen mass was either unchanged or slightly increased by AR-42 alone or in combination with GTx-024. As a gross measure of the systemic effects of treatment on immune function, these spleen mass results suggest AR-42 is not generally immunosuppressive and its activity is distinct from inhibitors of the JAK/STAT pathway in this context [28]. Unlike in gastrocnemius tissue, AR-42 treatment did not significantly suppress pSTAT3 signaling within the C-26 tumors (Figure 4F). Taken together, these multiple lines of evidence suggest that the anti-cachectic efficacy of AR-42 involves the inhibition of the IL-6/GP130/STAT3 axis in skeletal muscle tissue, but not systemic suppression of IL-6 or general immune signaling.

### Transcriptomic analyses of AR-42’s anti-cachectic effects in skeletal muscle

To further characterize AR-42’s anti-cachectic effects at the reduced dose of 10 mg/kg, RNA-seq analyses were performed on all gastrocnemius tissues from Study 1 (Figure 2A, B). This resulted in 31 evaluable samples across treatment groups (Supplementary Figure S8) after removal of two samples due to insufficient sequencing yield/quality. We detected 4,579 differentially expressed genes (DEGs; FDR < 0.1) in cachectic versus control muscle, whereas treatment of cachectic mice with GTx-024 or AR-42 alone resulted in 5,561 and 723 DEGs, respectively, consistent with their corresponding anti-cachectic efficacies (Figure 5A, Supplementary Figure S9A and B). Given the ability of HDAC inhibitors and androgens to modulate transcription, initial functional analyses were focused on curated *Mus musculus* transcription factor (TF) target gene sets, and revealed multiple over-represented TF targets in cachectic versus control muscle (Figure 5B). STAT3 and activation of transcription-1 (ATF1) gene sets were each represented twice in the top ten pathways following GSEA supporting their potential relevance in cachectic signaling. The two STAT3 target gene sets were combined and GSEA was repeated with the combined set for all treatment groups. In contrast to pSTAT3 activation (Figure 4D), this analysis demonstrated the inability of any treatment in tumor-bearing mice to significantly limit the importance of STAT3 target-gene regulation relative to cachectic controls (Figure 5C and Supplementary Figure S10). However, when analysis is focused on individual genes within the combined set that are differentially expressed in at least one comparison, clear cachexia-dependent regulation is apparent that responds only to AR-42 treatment (Figure 5D). A similar analysis with combined ATF-1 data sets revealed the ability of AR-42, but not GTx-024 treatment, to significantly impact ATF-1 target gene regulation in tumor-bearing mice implicating AR-42’s ability to modulate ATF-1 activation in its anti-cachectic efficacy (Figure 5E and Supplementary Figure S11). Of note, STAT3 and CEPBδ are among the differentially expressed ATF-1 target genes induced by cachexia that respond only to AR-42 treatment (Figure 5F).

**Figure 5.**
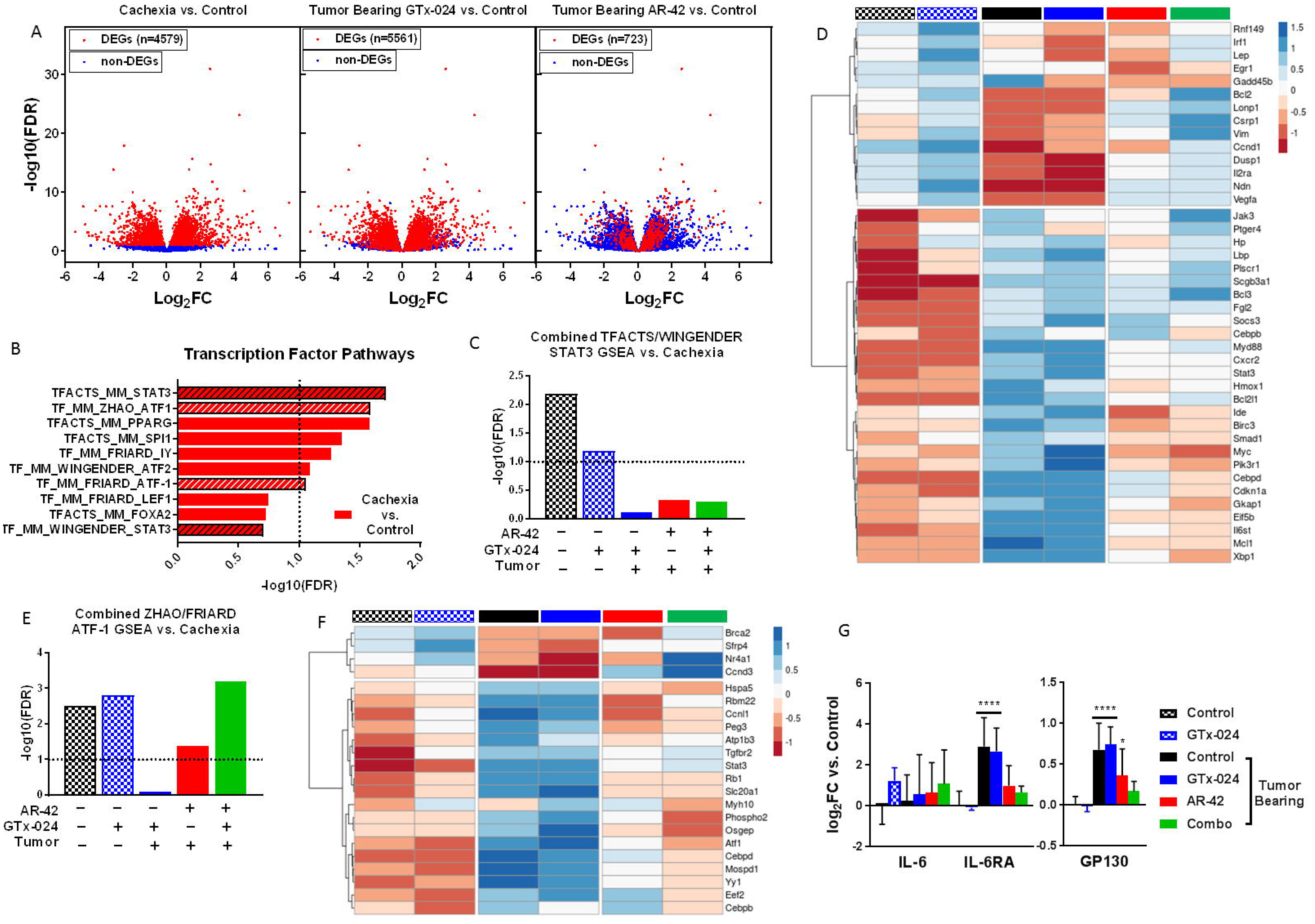
**A)** Effect of GTx-024 and AR-42 monotherapies on cachexia-related differentially regulated genes (DEGs) from RNA-seq analyses of Study 1 gastrocnemius muscles. All three panels consist of individual genes plotted with respect to their log2 fold change and −log10 Benjamini-Hochburg adjusted p-values from the comparison of cachexia vs tumor-free controls. Colors of the points reflect the DEG status of each gene for the given comparison. **B)** Results from transcription factor pathway-focused GSEA of tumor-bearing (cachexia) versus tumor-free control transcriptomes. STAT3 and ATF-1 gene sets used for subsequent combined analyses are hatched. **C)** Significance values from GSEA using combined STAT3 gene sets identified in **B.** Each treatment group is compared to tumor-bearing control (cachexia) transcriptomes. **D)** Heat map of DEGs within the combined STAT3 gene sets representing mean z scores calculated from normalized RNAseq count data. Tumor-free control (black checkered), GTx-024-treated tumor-free (blue checkered), tumor-bearing control (black), GTx-024-treated tumor-bearing (blue), AR-42-treated tumor-bearing (red) and Combination-treated tumor-bearing (green). **E)** Results of GSEA using combined ATF-1 gene sets identified in **B** across treatment groups versus tumor-bearing control (cachexia) transcriptomes. **F)** Heat map of DEGs within the combined ATF-1 gene sets (mean z score). Treatment groups are as in **D. G)** mRNA expression of mediators of IL-6 signaling upstream of STAT3. Data presented as mean±SD of per animal log-transformed fold change (log_2_FC) values versus tumor-free controls. *p<0.1, **p<0.05, ***p<0.01, ****p<0.001 based on Benjamini-Hochburg adjusted p-values from DESeq2.

We further evaluated the expression of genes within the IL-6 pathway as IL-6-mediated STAT3 target gene regulation is well characterized in the C-26 model [17], and IL-6-mediated increases in skeletal muscle cyclic AMP (cAMP), a primary driver of ATF-1 activation [29], have also been reported[30]. Unlike circulating IL-6 cytokine, IL-6 mRNA in gastrocnemius muscle was not induced by cachexia, nor was it modulated by any treatment (Figure 5G). However, expression of both IL-6 receptor (IL-6RA) and the key effector GP130 were elevated in cachectic mice and required AR-42 (IL-6RA) or combination treatment (GP130) to restore to non-cachectic control levels.

Considerable overlap exists between the transcriptomes of cachectic gastrocnemius muscles from mice treated with 10 or 50 mg/kg AR-42 such that high fold-change DEGs identified by Tseng *et al.* [17] and in the current study are all regulated in the same direction (n= 209, Supplementary Figure 12A-B). Similar to previous analyses (Supplementary Figure S5), functional interrogation of the genes within this overlap further support the importance of AR-42’s ability to modulate immune and extracellular matrix signaling in eliciting its anti-cachectic effects (Supplementary Figures S12C). Taken together these findings support the ability of the reduced 10 mg/kg dose of AR-42 to generate anti-cachectic effects by reducing procachectic IL-6RA/GP130/STAT3 signaling in skeletal muscle.

### Transcriptomic analyses of GTx-024’s anabolic effects in skeletal muscle

To better understand GTx-024’s contribution to the efficacy apparent in combination-treated mice, the transcriptome of combination-treated gastrocnemius muscle was compared to cachectic controls revealing 2,026 DEGs (or 50.6% of all DEGs) not solely attributable to AR-42 treatment (Figure 6A). We hypothesized that GTx-024-mediated anabolic signaling detectable in GTx-024-treated tumor-free controls would be diminished in tumor-bearing GTx-024-treated animals in the absence of AR-42. Though very few DEGs were apparent in GTx-024-treated tumor-free controls (n = 27, Supplementary Figure S13), GSEA focused on TF pathways revealed abundant coordinated signaling with regulation of β-catenin (CTNNB1) target genes providing the most significant overlap (FDR < 1e-5, Figure 6B). Coordinate regulation of β-catenin target genes was not apparent in cachectic controls or following GTx-024 or AR-42 monotherapy, but was again among the most prominent pathways detected by GSEA in combination-treated mice (FDR < 1e-5, Figure 6C). GSEA plots demonstrate a robust pattern of GTx-024-mediated activation of β-catenin target genes requiring AR-42 co-administration in cachectic mice (Figure 6D, *leftmost panel compared to rightmost panel*). Analysis of overlap of the leading edge genes revealed a large number of CTNNB1 target genes regulated by both GTx-024 and cachexia versus tumor free controls but in different directions (n=49 *middle*, 17 *bottom left*; Supplementary Figure S14A). Many fewer leading edge genes were regulated by AR-42 monotherapy but also in an opposite direction to GTx-024 (n= 2 *middle*, 23 *top middle*; Supplementary Figure S14B). However, combined therapy results in a larger leading edge gene set overlap that is regulated in a similar direction to GTx-024 monotherapy (n= 29 *middle*, 37 of 47 *top middle*; Supplementary Figure S14C). This pattern of β-catenin target gene regulation is also apparent when DEG’s within the TFACTS_CTNNB1 gene set are visualized across treatment groups (Supplementary Figure S15).

**Figure 6.**
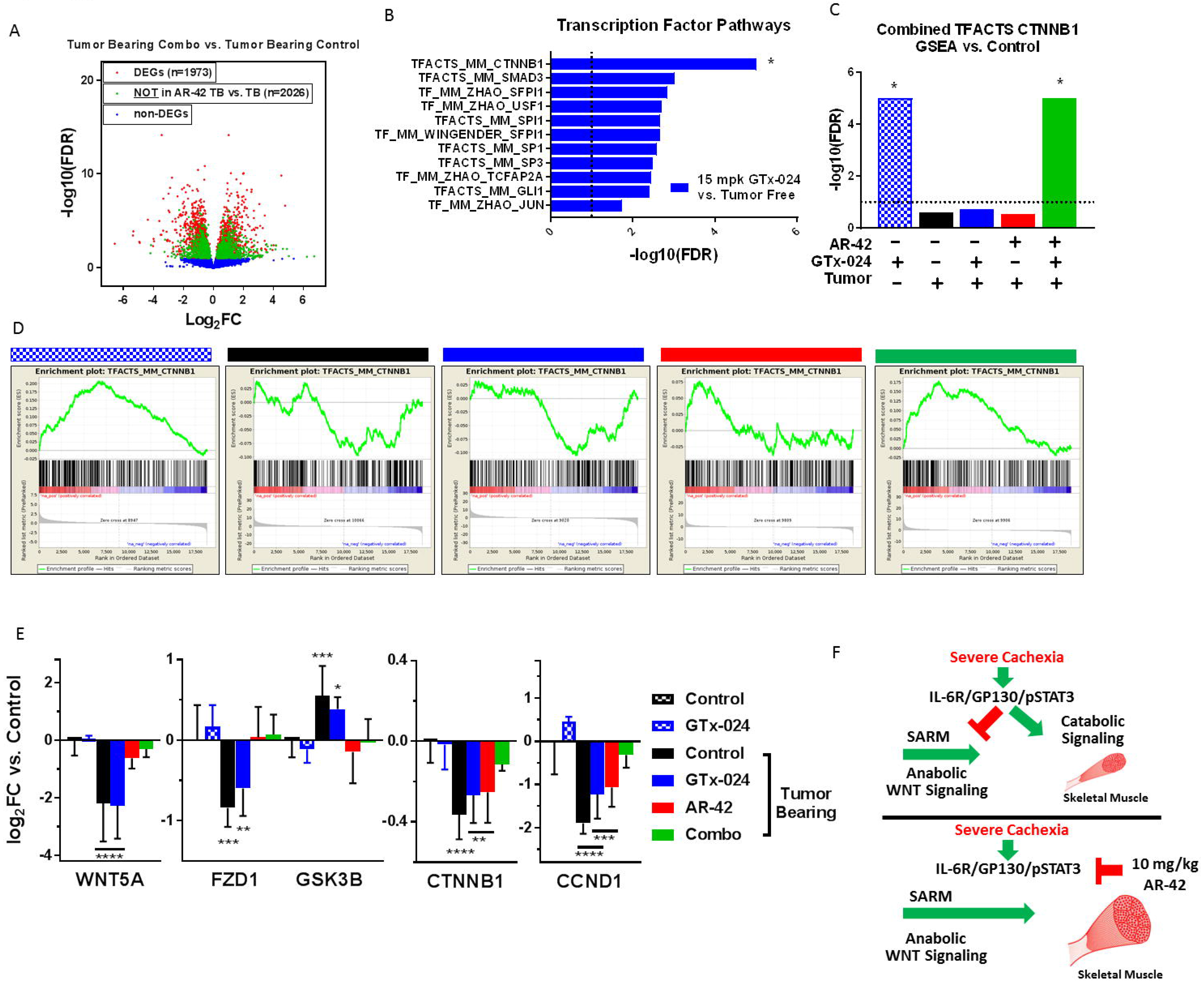
**A)** Volcano plot from RNA-seq analyses of Study 1 gastrocnemius muscles for tumor-bearing combination-treated mice versus tumor-bearing controls. Genes not differentially expressed in this comparison are indicated with blue. The remaining genes (red and green) are DEGs in the combination-treated versus tumor-bearing control comparison. The green coloring indicates the subset of these DEGs that are not also differentially expressed in the comparison of AR-42-treated tumor-bearing mice versus tumor-bearing controls, suggesting these genes are responsive to only the combination therapy. Log2-transformed fold change (FC) in expression is plotted on the x-axis and −log10 transformed Benjamini-Hochburg adjusted p-values are plotted on the y-axis. **B)** Significance values from transcription factor pathway-focused GSEA of GTx-024-treated tumor-free versus tumor-free control transcriptomes. *FDR< 1e-5 was determined for the CTNNB1 gene set, and was set to 1e-5 for the plot. **C)** Significance values from GSEA using combined CTNNB1 gene sets. Each treatment group was compared to tumor-free control transcriptomes. *FDR < 1e-5 determined, set to 1e-5 for plot. **D)** Enrichment plots from GSEA of the CTNNB1 gene set for each treatment group versus tumor-free control comparisons. GTx-024-treated tumor-free (blue checkered), tumor-bearing control (black), GTx-024-treated tumor-bearing (blue), AR-42-treated tumor-bearing (red) and Combination-treated tumor-bearing (green). **E)** mRNA expression of WNT effectors upstream of β-catenin. Data are presented as mean±SD of log-transformed fold change (log2FC) values versus tumor-free controls. *p<0.1, **p<0.05, ***p<0.01, ****p<0.001 based on Benjamini-Hochburg adjusted p-values from DESeq2 **F)** Graphical Mechanistic Hypothesis.

Expression of the canonical skeletal muscle WNT agonist Wnt5a (WNT5A), canonical WNT receptor Fzd1 (FZD1) and β-catenin itself (CTNNB1) were all reduced with C26 tumor burden, whereas the negative regulator of β-catenin, GSK3B, was up-regulated (Figure 6E). In each case, GTx-024 monotherapy in tumor-bearing mice failed to restore expression to tumor-free control levels. However, with the exception of β-catenin, AR-42 treatment effectively reversed tumor-induced regulation. Furthermore, combination treatment alone restored β-catenin and the well-characterized β-catenin target gene cyclin D1 (CCND1) [31] expression to tumor-free control levels. Taken together these data provide strong support for: 1) the dependence of GTx-024’s anabolic effects in skeletal muscle on functional WNT/β-catenin signaling; 2) C-26 tumor burden’s ability to disrupt WNT/β-catenin signaling in skeletal muscle; and 3) AR-42’s ability to restore WNT/β-catenin responsiveness to treatment with GTx-024.

## DISCUSSION

### Anti-cachectic efficacy of reduced dose 10 mg/kg AR-42

AR-42 is currently under clinical evaluation as a direct anti-tumor agent (NCT02282917, NCT02795819, NCT02569320). Recent clinical experience suggested the previously described 50 mg/kg anti-cachectic dose in mice would be poorly tolerated and inconsistent with administration to already heavily treated cachectic cancer patients. We describe a 5-fold AR-42 dose reduction that retained anti-cachectic efficacy across multiple studies (Figure 2 and Supplementary Figure S4). Preliminary enrichment analyses of AR-42-regulated transcripts in muscle implicated IL-6 and immune system signaling in the anti-cachectic efficacy of AR-42 (Supplementary Figure S6). However, at this reduced dose of AR-42, circulating cytokines were not significantly modulated by AR-42 treatment (Figure 4A and Supplementary Table S1). By contrast, activation of STAT3, an essential mediator of IL-6 family cytokine-derived immune signals, was AR-42-sensitive in skeletal muscle (Figure 4D). Seto *et al*. have recently demonstrated that IL-6 family cytokine signaling through STAT3-dependent, as opposed to FOXO-, NF-κB-, SMAD- or C/EBP-dependent transcription, drives C2C12 myotube atrophy in response to C-26 cell conditioned media [32]. In agreement with the critical role of STAT3 activation in C-26-mediated cachexia, both genetic manipulation and pharmacological inhibition of STAT3 mitigate C-26 tumor-induced losses in skeletal muscle [24, 32]. Transcriptomic analyses of gastrocnemius muscle confirmed the ability of reduced dose AR-42 to markedly impact cachexia-associated transcription (Figure 5A) and substantiated STAT3 and ATF-1 transcriptional programs as cachectic drivers (Figure 5B). ATF-1 is a member of the cAMP response element–binding protein (CREB) family of TFs whose activation is associated with fibroblast proliferation and transformation, but has no described role in muscle wasting [33].

AR-42 treatment reduced the mRNA expression of IL-6RA and the effector protein GP130 (Figure 5G) similar to reports of AR-42 activity in multiple myeloma cells [26] and the activity of pan-HDACi’s in naïve CD4+ T cells [34]. Tissue-specific HDACi-mediated muting of IL-6R and/or GP130 induction following cachectic challenge provides a plausible mechanism for the reversal of IL-6 family cytokine-driven ATF-1/STAT3 transcription (Figure 5D and 5F) we detected in the absence of broader systemic immune effects (Figure 4). Determining the precise mechanism by which AR-42 treatment, but not treatment with other HDACi’s[15], mediates anti-cachectic efficacy will require further study, but our data support the continued evaluation of AR-42 as a compelling anti-cachectic agent.

### Impact of C-26 tumor burden on androgen signaling

A key finding of this report is the extent of resistance to anabolic androgen administration in the C26-model of cancer cachexia. We utilized fully anabolic doses of two SARMS and a potent steroidal androgen, administered orally (GTx-024) and parenterally (TFM-4AS-1, DHT), which resulted in no detectable anti-cachectic efficacy (Figure 2), despite demonstrated anabolic capability (Supplementary Figure S2) and evidence of systemic hormonal activity (Supplementary Figure S3A). Our results reflect the dearth of published reports of anabolic therapeutic efficacy in this common model. At the time of manuscript preparation, we identified a single demonstration of anti-cachectic anabolic therapy in C-26 mice, despite anabolic agents representing the most advanced clinical development programs in cancer wasting [35].

Androgens have a well characterized ability to normalize skeletal muscle catabolic gene expression associated with either glucocorticoid (dexamethasone)- or hypogonadism (castration)-induced atrophy [13, 36, 37]. We hypothesized that the inability of androgens to reverse C-26 tumor-mediated atrogene expression underlies their lack of efficacy (Figure 3A). Consistent with this hypothesis, inflammatory cytokine-driven catabolic signaling in the C-26-model, which is mechanistically distinct from androgen-responsive wasting, appears completely insensitive to androgen administration (Figure 5). A plausible explanation for androgens’ ineffectiveness as a monotherapy is a cachexia-mediated direct disruption of AR signaling.

However, the response of the hypothalamic-pituitary-gonadal axis (Supplementary Figure S3A), several cytokines (Figure 4A) and gastrocnemius transcriptome (Supplementary Figure S13) to androgen, along with no obvious effects of tumor burden on AR mRNA or protein (Figure 3B and C), suggests the AR’s ability to respond to androgen in skeletal muscle remains intact. Nonetheless, catabolic signaling through the IL-6/GP130/STAT3 axis appears refractory to diverse androgen administration.

In addition to mitigating catabolic proteasomal signaling, androgens have well-characterized direct anabolic effects on skeletal muscle that include targeting MUSCs and pluripotent mesenchymal progenitor cells to promote muscle hypertrophy [12]. We hypothesized that compromised androgen-mediated anabolic signaling might contribute to GTx-024’s lack of anti-cachectic efficacy. For consistency across studies, all of our mechanistic analyses focused on gastrocnemius muscle which, like most skeletal muscles, has scant AR expression (Figure 3C), but readily responds to androgen administration [37]. As such, GTx-024 treatment in tumor-free mice resulted in very few DEGs (Supplementary Figure S13), but GSEA, which is designed to detect patterns within whole transcriptomes, as opposed to individual DEGs [38], revealed a robust induction of β-catenin target gene regulation (Figure 6B-D). Our results are consistent with androgen-mediated β-catenin activation reported in the context of whole muscle tissue [39], and as a requirement for myogenic differentiation of pluripotent mesenchymal cells [40]. Notably, GTx-024-mediated β-catenin target gene regulation is completely abrogated in the context of C26-tumor burden (Figure 6C-D), which corresponds with coordinated suppression of canonical WNT pathway effectors (Figure 6E). GTx-024-mediated β-catenin activation was only restored in the presence of AR-42 which, as a monotherapy, normalized WNT effector expression. Elucidating AR-42-responsive cachectic signals governing WNT suppression warrants further interrogation.

To the best of our knowledge, this is the first report of dysfunctional skeletal muscle WNT signaling in experimental cachexia. Cachexia was associated with suppression of canonical WNT effectors (Figure 6E) and an inability to respond to androgen-mediated WNT signals (Figure 6D). Multiple β-catenin target genes did respond to cachectic signaling (Supplementary Figure S14 and S15), suggesting that components of WNT-mediated β-catenin target gene regulation remain intact despite the suppression of upstream WNT effectors. Importantly, both constitutive activation and genetic abrogation of WNT signaling impair proper adult MUSC function in response to injury [41–43]. Our data suggest tightly controlled WNT-signaling is lost in tumor-bearing mice. This is consistent with other reports of MUSC dysfunction in the C-26 model [44]. Intriguingly, β-catenin-mediated follistatin induction is required to promote MUSC differentiation following stimulation with WNT ligands[45] and androgens [46]. Given the clear effects of exogenous androgen administration on MUSC activation [47], it is plausible that cachexia-mediated disruption of WNT signaling represents a functional blockade of androgenic anabolism in skeletal muscle (Figure 6G). Furthermore, intact WNT signaling is required for proper MUSC function irrespective of androgen administration, suggesting the dysfunctional WNT signaling reported here might be linked more broadly to the important clinical problem of cancer-induced anabolic resistance [48].

We recognize that our experimental paradigm is limited in a number of ways. We have evaluated a single rapid model of experimental cachexia, which necessarily limits the broader interpretation of our findings. The short treatment window (<14 days) afforded by the C-26 model in our hands also severely curtailed our ability to demonstrate overt anabolic effects following GTx-024 treatment relative to other anabolic agents in less severe models of cachexia [49]. Furthermore, our studies were limited to fixed dose levels. Ten mg/kg AR-42 was the lowest effective dose evaluated as monotherapy, but its ability to augment anabolic therapy at even lower doses was not investigated. Likewise, the dose of GTx-024 employed was likely much higher than what is required to provide maximal anabolic benefit. Further dose optimization would be critical to both improve efficacy and minimize toxicity as the tolerance for additional side effects ascribed to anti-cachexia therapy in already heavily treated cancer patients is low.

### Combined anabolic and anti-catabolic therapy in cancer cachexia

To our knowledge, this is the first report combining SARM and HDAC inhibitor administration in experimental cachexia, which demonstrated efficacy using two agents currently undergoing clinical development. Even with the limited treatment window available in the C-26 model, we show improved total body weight (Figure 2A, D), lower limb skeletal muscle mass (Figure 2B, E), and grip strength (Figure 2C, F) for two different SARMs when combined with AR-42 over tumor-bearing controls and SARM monotherapy. Transcriptome characterization of skeletal muscle tissue revealed the ability of AR-42, but not GTx-024, to ameliorate IL-6/GP130/STAT3-mediated catabolic signaling, whereas GTx-024, but not AR-42, stimulated anabolic canonical WNT signaling. Strikingly, GTx-024’s ability to effectively stimulate WNT signaling required AR-42 co-treatment in cachectic mice. Notably, when AR-42 was combined with DHT, terminal body weights were significantly improved compared to single agent AR-42 treatment (Figure 2D). Our mechanistic support for beneficial signaling in muscle following SARM and HDACi co-administration along with DHT’s *in vivo* efficacy suggests that similar results are possible with optimized combination SARM regimens.

Despite established efficacy in diverse patient populations [50, 51], GTx-024 failed to provide anabolic benefit in advanced NSCLC patients [52]. Though weight loss was not required for enrollment in GTx-024’s registration trials, roughly half of all patients reported >5% unexplained weight loss at initiation of chemotherapy suggesting a high prevalence of cachexia at diagnosis. In a similar cohort receiving anabolic ghrelin mimetic anamorelin therapy, subgroup analyses revealed patients with body mass indices <18.5 (and presumably severe cachexia) showed no improvements in body composition [53]. Analogous to these clinical populations, our data show that anabolic androgen administration cannot overcome severe catabolic signaling in the C-26 model and that profound cachectic burden additionally results in a blockade of critical anabolic signaling. Furthermore, we show that AR-42’s anti-cachectic efficacy involves both mitigating catabolic signaling and licensing anabolic signaling providing compelling mechanistic support for combined GTx-024/AR-42 administration in cachectic patients. Combination therapy demonstrates the potential to improve anabolic response in patient populations with advanced cancer wasting.

## ACKNOWLEDGEMENTS

We thank Arno Therapeutics, Inc., for generously providing AR-42. We also thank the Molecular Carcinogenesis and Chemoprevention Program (OSU Comprehensive Cancer Center), as well as Jiang Wang and the Pharmacoanalytical Shared Resource and Genomic Shared Resource in The Ohio State University Comprehensive Cancer Center which is supported by NCI/NIH Grant P30-CA016058. We are grateful to Dr. Appaso Jadhav and Ms. Uma Subrayan (The Ohio State University College of Pharmacy) for verification of GTx-024 purity and technical assistance in preparation of tissue samples for analysis. The LH assays were performed by The University of Virginia Center for Research in Reproduction Ligand Assay and Analysis Core which is supported by the Eunice Kennedy Shriver NICHD/NIH (NCTRI) Grant P50-HD28934. This work was also supported in part by NCI/NIH K12-CA133250-07 (Dr. Coss), Eli Lilly Fellowship (S. Liva) and Pelotonia Idea Award (OSU Comprehensive Cancer Center, Dr. Coss).

